# The structural and functional connectivity neural underpinnings of body image

**DOI:** 10.1101/2020.08.04.236547

**Authors:** Massieh Moayedi, Nasim Noroozbahari, Georgia Hadjis, Kristy Themelis, Tim V. Salomons, Roger Newport, Jennifer Lewis

## Abstract

How we perceive our bodies is fundamental to our self-consciousness and our experience in the world. There are two types interrelated internal body representations—a subjective experience of the position of a limb in space (body schema) and the subjective experience of the shape and size of the limb (body image). Body schema has been extensively studied, but there is no evidence of the brain structure and network dynamics underpinning body image. Here, we provide the first evidence for the extrastriate body area (EBA), a multisensory brain area, as the structural and functional neural substrate for body shape and size. We performed a multisensory finger-stretch illusion that elongated the index finger. EBA volume and functional connectivity to the posterior parietal cortex are both related to the participants’ susceptibility to the illusion. Taken together, these data suggest that EBA structure and connectivity encode body representation and body perception disturbances.

## Introduction

The representation of our body is essential for how we interact with our environment. These representations arise from multimodal sensory inputs, including visual, tactile, proprioceptive, interoceptive, nociceptive, and motoric inputs [1]. There are at least two proposed implicit body representations: the body schema, which encodes the position of the body in space, and the body image, which refers to the subjective experience of the size, shape, and features of the body [1,2]. The structural and functional neural network representation of body image has yet to be elucidated.

Illusion studies are a reliable method to disturb body image under controlled conditions and are therefore powerful tools to investigate the neural mechanisms underlying body image [3]. For example, the Pinocchio illusion, and various derivatives of this unimodal illusion, have been used to induce body perception disturbances in healthy individuals [4,5]. The illusory disruption of the body image can be reinforced with cross-modal illusions, which simultaneously manipulate two or more sensory channels [6]. For example, the addition of a tactile stimulus can reinforce a visual illusion and subsequently enhance the robustness of the illusion, and thus susceptibility to the illusion. Integrating tactile and visual information requires multisensory processing and binding in higher order cortical regions. Indeed, brain imaging studies of visuo-tactile illusions have identified the bilateral ventral premotor cortex (PMv), left posterior parietal cortex (PPC), and occipitotemporal areas [7–12] to be implicated in body representation. Unlike these previous illusion studies that modify body perception by targeting body schema, here, we specifically and exclusively modify body image.

The finger-stretch illusion is a robust visuo-tactile illusion in which participants experience alterations of their index finger in a computer-mediated augmented reality system with congruent sensory feedback from the experimenter [13]; *i.e.* the participant’s finger appears to be elongating while the experimenter pulls on the tip of the finger, and shortening back to actual size while the experimenter pushes on the tip of the finger. An advantage of the finger-stretch illusion is that, unlike other visuo-tactile illusions (*e.g.* the rubber hand illusion), the finger-stretch illusion is applied to the participant’s own body through congruent manipulation of visual, proprioceptive and tactile stimuli [14], rather than incorporating a non-body object. Here, we aim to use the finger-stretch illusion to identify the structural and functional network underpinnings of changes to body image representation. Here, we provide the first evidence for the extrastriate body area (EBA), a multisensory brain area, as the structural and functional neural substrate for body shape and size.

## Material and Methods

This study comprised two independent investigations: a behavioural study to determine whether and distal versus a proximal perspective affects the effectiveness and features of the body illusion; and second, an imaging study to identify the neural correlates of the illusion.

### Participants

#### Behavioural study (Experiment 1)

Twelve right-handed adults (7 women, 5 men), aged (mean ±SD) 23.6 ±2.3 years were recruited for this study from the University of Nottingham, Nottingham, UK. Participants provided written informed consent to procedures reviewed and approved by the University of Nottingham ethics committee in line with the Declaration of Helsinki.

#### Imaging study (Experiment 2)

Twenty healthy adults were recruited from the University of Reading, Berkshire, UK. Participants provided written informed consent to procedures reviewed and approved by the procedures reviewed and approved by the University of Nottingham ethics committee in line with the Declaration of Helsinki in line with the Declaration of Helsinki, and were compensated for their time. Nineteen participants were scanned as one was excluded due to MRI contraindications. A further participant’s dataset was excluded due to technical problems during the scan. Therefore, of the final sample of 18 participants (10 women, 8 men), aged 24.3 ±5.9 years, 17 were right-handed, and one was left-handed

### MRI data acquisition

Functional brain images were collected on a 3T MRI scanner (Magnetom Trio; Siemens, Erlangen, Germany) using a 32-channel head coil. For each participant, four runs of a 162 whole brain image timeseries (5 min 24 s) were obtained using a gradient-echo, echo-planar scanning sequence (repetition time (TR)=2 s, echo time (TE)=29 ms, flip angle=90°, GRAPPA=2, field of view=272 mm^2^, 30 axial slices, slice thickness 3.5 mm, no gap, voxel size=2.1 × 2.1 × 3.5 mm^3^). A high-resolution anatomical scan was also acquired (T1-weighted, 3-dimensional magnetization-prepared rapid acquisition gradient echo sequence scan, TR=2s, TE= 2.99ms, FOV=250mm, 192 sagittal slices, GRAPPA=2, voxel size = 1.0 × 1.0 × 1.0 mm^3^).

### Experimental design

#### Behavioural study

We first sought to determine whether observing a finger-stretch illusion in an MRI scanner—i.e. a distal setup rather than a proximal one—would affect the illusory experience. Participants underwent a finger-stretch illusion under two conditions (Fig 1):(1) proximal and (2) distal setup. In setup 1, seated participants positioned their hand within the MIRAGE device, a Mediated Virtual Reality (MVR) system (University of Nottingham, Nottingham, UK). The Mirage device uses a camera and mirror arrangement where one can view real-time video images of their hand in the same location as their actual hand. Instantaneous digital manipulations to the visual input give rise to a range of bodily illusions. We performed a finger stretch on the index finger of the left hand. This decision was based on the physical constraints in the MRI environment. Setup 2 was identical to setup 1 with one key difference: the video image of the hand was on a screen two meters in front of the participant, thus removing the egocentric aspect of the illusion. This mimicked the setup inside the MRI scanner. Participants received two stretches per condition. Both setups used the same illusion on the participants’ left index finger. Participants rated six statements about each stretch on a 11-point Likert scale, anchored at “not at all” and “extremely.” The statements included two control statements: 1. ‘*It felt like I had 2 left index fingers,*’ 2. ‘*My finger was getting hot*,’; and four illusion susceptibility statements: 1.’*I feel like the finger I’m seeing belongs to me*’ 2. ‘*I feel like I’m watching myself’* 3. ‘*I feel like my finger is longer than normal*’, 4. ‘*It felt like my finger was really being stretched.’* (Fig 1c).

**Fig 1.**
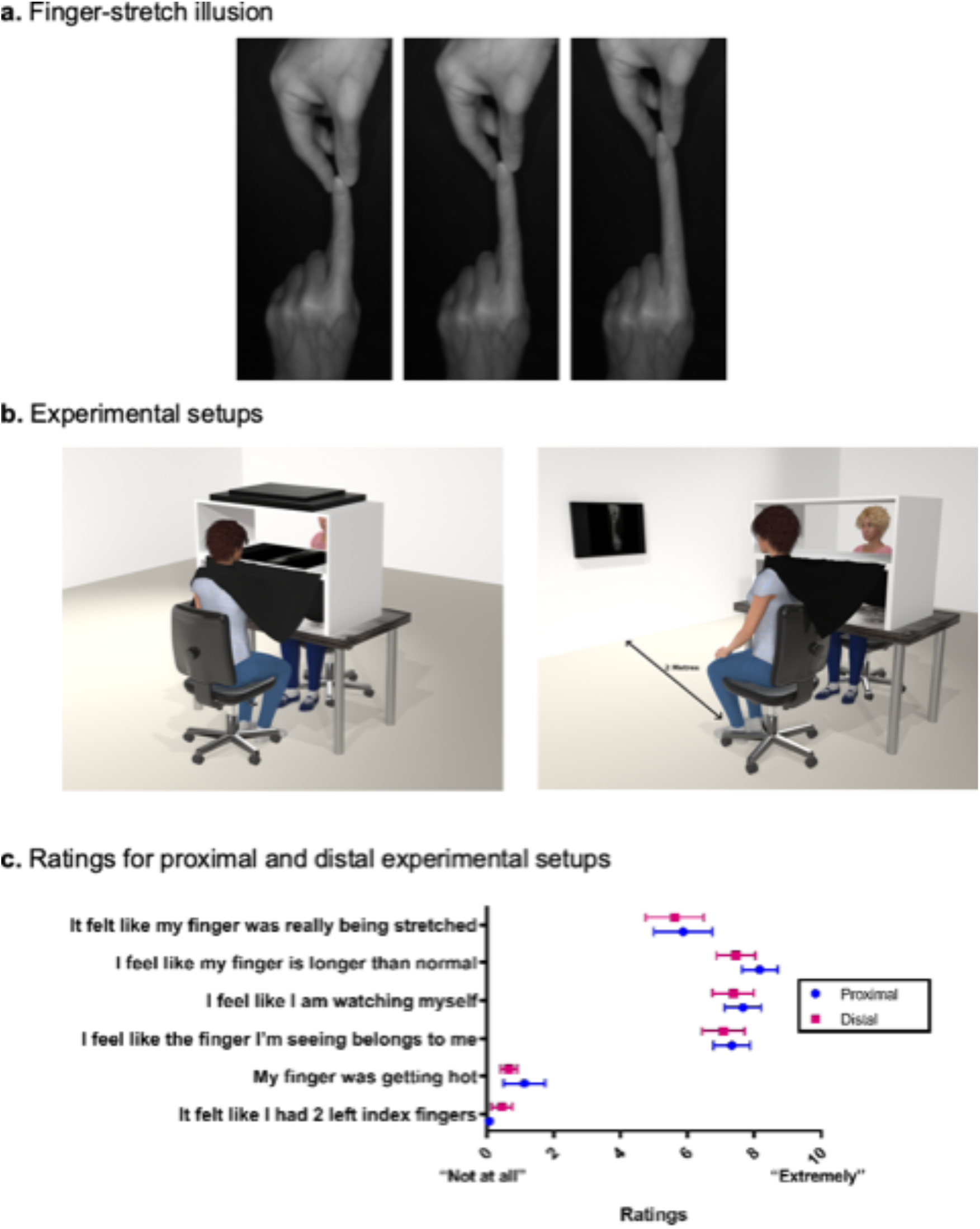
Behavioural experiment to determine whether the finger-stretch illusion can be performed within an MRI scanner with a different egocentric setup. (a). The finger-stretch illusion is a visuo-tactile illusion in which participants experience their index finger elongating in a computer-mediated augmented reality system with congruent sensory feedback from the experimenter. The progression of the illusion is shown from the left panel to the right. As the image of the finger is elongated, the experimenter pulls on the tip to add tactile feedback. (b). The illusion was tested in two experimental setups: (1) proximal (which is the original setup) and (2) distal (n=12). The distal setup is similar to that which will be performed in the MRI. In the distal setup, the participant is watching a digital image of their finger undergoing the finger-stretch illusion. The screen is 2 meters away from the participant. (c). Mean and standard error of the mean ratings of distal (red) and proximal (blue) setup are depicted. There were no significant differences between ratings for each of the setups, indicating that the feeling that the participant’s own finger was stretched is similar and equally effective in the distal as the proximal setup (One-way repeated-measures ANOVA, *F*_1,11_= 1.433, *p*= 0.256, all post-hoc t-tests *p*> 0.05).

#### Imaging study

All participants were naïve to the MIRAGE finger-stretch illusion [13,14]. As participants lay supine in the scanner, an MR-compatible camera in the MR-environment captured real-time digital images of the left index finger, to a computer in the control room. Participants viewed the image of the left index finger on a 32” fMRI compatible LCD screen (BOLDScreen, Cambridge Research Systems, Rochester, UK) through a mirror mounted onto the head coil. The experimenter stood next to the MRI table, holding the participant’s left index throughout the MRI scan. The participants’ remaining fingers were covered with a black, non-reflective cloth to be invisible in the image. The MIRAGE finger stretching illusion was run in the LabView Software package v 15.0 (National Instruments, Austin, TX, United States of America) on an Apple MacBook Pro (Retina, 13-inch, Early 2015, Apple, Cupertino, CA, United States of America), running Windows 2008 (Microsoft, Seattle, WA, United States of America). The illusion (‘manipulation’ condition) comprised of three distinct phases: index finger elongation (Fig 1a), maintenance of finger elongation, and shrinking the finger back to normal size. During finger elongation, participants observed their finger being lengthened while the experimenter simultaneously pulled gently on the distal tip of their finger. During maintenance, participants observed the experimenter holding the tip of the lengthened index finger. During shrinking, participants observed their index finger shrinking back to normal size, while the experimenter simultaneously pushed the tip of their finger. The experimenter was cued by an auditory signal on which tasks to perform. Each phase was 3 seconds, and the whole illusion cycle took 9 seconds. We also performed a control condition (non-manipulation), where the same three illusion phases occurred, but without the finger being visually lengthened. Manipulations were identical across the group, and both participants and experimenters were blinded to the condition. During a 10s period, eight seconds (s) after each illusion, participants rated the statement: “*I felt like my finger was really being stretched*” on a 6-point rating scale, where 0 represented ‘not at all’ and 5 represented ‘extremely lengthened’. The next illusion occurred 12s after rating, with a pseudo-randomized jitter of 0-3 s, (rectangular distribution). The experiment utilized a block design, with four trials each of illusion and control conditions presented in a pseudo-randomized and counterbalanced order across four runs, for a total of 16 trials per condition.

### Data preprocessing

#### Ratings

Each participant rated each trial of the finger-stretch and the control conditions. These ratings were averaged across all trials for each illusion. Susceptibility scores were calculated by subtracting the control ratings from the finger stretch illusion ratings.

#### Voxel-based Morphometry

To examine grey matter correlates of the susceptibility scores, we performed voxel-based morphometry (VBM)44 in the Statistical Parametric Mapping (v12; (http://www.fil.ion.ucl.ac.uk/spm/software/spm) DARTEL toolbox [45]. Briefly, preprocessing included setting the origin of the image at the anterior commissure of each subject, affine spatial normalization, tissue segmentation. Next the various tissue classes were meaned, and were then aligned to create a template. Deformations from this template to each of the individual images were computed, and the template was then re-generated by applying the inverses of the deformations to the images and averaging. This procedure was repeated several times. The template was then normalized to MNI152 space. Finally, warped images were generated, and spatially smoothed with an 8 mm FWHM Gaussian kernel.

#### Illusion Task Functional MRI analysis

All imaging analysis was performed using FSL (FMRIB’s Software Library, v.5.0, Oxford, UK) [46], unless otherwise indicated. Prior to statistical analysis, non-brain structures were removed from each participant’s structural images by Brain Extraction Tool (BET v.2.0). Preprocessing steps were performed using the Multivariate Exploratory Linear Optimized Decomposition into Independent Components (MELODIC) [47] toolbox. The first 5 volumes were removed for each participant to allow for signal equilibrium, and a high pass filter cut off of 100 s (0.01 Hz) was applied. Standard preprocessing, including motion correction using MCFLIRT [48], slice timing correction, spatial smoothing with a 5 mm (FWHM) Gaussian kernel was applied. The functional images were registered to the Montreal Neurological Institute (MNI)-152 template. Data were then ICA-denoised by two independently trained raters (NN and AY). To further denoise the dataset, we performed aCompCor procedures [49]. Briefly, signals from the cerebrospinal fluid (CSF) and white matter (WM) were extracted. The CSF and WM masks were 5mm spheres manually drawn in FSLview on the MNI-152 2-mm brain image. Principal components analysis (PCA) was performed on each of the timecourses in Matlab v9.4 (Mathworks, Natick, MA, USA), and the first 5 components were regressed out of the fMRI data for each participant.

### Statistical Analysis

#### Behavioural Study

A 2-way repeated-measures ANOVA was used to test significant differences between the two illusion setups. Post-hoc paired t-tests were performed to identify significant differences in ratings between proximal and distal setups. Significance was set at *p*<0.05.

#### Voxel Based Morphometry

A whole brain, voxelwise statistical analysis was performed using the general linear model to determine which grey matter regions correlated with susceptibility scores. The model included a regressor for group (to model the intercept), and demeaned susceptibility scores. Statistical images were thresholded at a cluster-corrected *p*_*FWE*_<0.05, with a cluster-forming height threshold p<0.001.

#### Illusion Task Functional MRI analysis

A general linear model (GLM) analysis was carried out on the preprocessed and denoised data using FEAT (FMRI Expert Analysis Tool) version 6.00 of FSL (FMRIB’s Software Library). Three different contrasts were modelled in the design matrix: finger-stretch, control, and finger-stretch>control. Motion parameters were not included in the model, as motion-related artefacts were corrected in preprocessing. A fixed-effect analysis was used to calculate a mean for each contrast across the four fMRI runs. Group-level analyses were performed using FLAME 1+2 (FMRIB’s Local Analysis of Mixed Effects) for each contrast. Statistical images were thresholded using a corrected-cluster p<0.05 (cluster-forming height threshold Z>3.1). We also performed a conjunction analysis to identify brain regions that were activated by both the finger-stretch and control illusions [50]. Briefly, the script identifies regions of significant overlap across two statistical images. Significance was set at a corrected-cluster p<0.05 (cluster-forming height threshold Z>3.1).

#### Psychophysiological Interaction Analysis

After identifying the regions of interest that displayed altered activation during the illusion task, a psychophysiological interaction (PPI) [51] analysis was performed to identify other voxels in the brain that displayed coupled activation with activity of these regions. Seed ROIs were based on peak functional activations from the initial group level FEAT analysis for illusion>control contrast (all coordinates are reported in MNI152 space (X,Y,Z): right PPC (50, 26, 42); left PPC (46, −40,54); right EBA (48, −64,4), Left EBA (−42,−74,−2), right FBA (47,−60,−11), left FBA (−46,−58,−16) and PMv (−46, 10, 30). ROIs were drawn using a 5-mm radius sphere centred at the peak voxel and each mask was transformed to individual functional space using the FSL linear registration tool (FLIRT) [48,52].

PPI analysis was initially performed on individual datasets by first extracting deconvolved MR signals from each seed ROI. This extracted time course represents an approximation of neural activity which was centered and multiplied by a psychological factor (task condition). The PPI regressor was determined by the interaction between the time course and the psychological regressor (onset of illusion). Both psychological and physiological regressors along with the PPI term were used in a GLM analysis.

Individual linear contrasts of PPI were subsequently grouped for analysis (FLAME 1 and 2, mixed effect). Statistical threshold was set at a corrected-cluster p<0.05 (cluster-forming height threshold *Z*>3.1). To restrict the search to regions that show functional connectivity with the EBA (−52,−64,−2) and the FBA (40,−52,−20), meta-analytic functional connectivity maps for each ROI were created using Neurosynth (www.neurosynth.org) [53]. These maps were used to restrict the search areas of their respective PPI during individual level analysis. The Neurosynth EBA functional connectivity map was thresholded at Z ≥ 5 and that of the FBA was thresholded at Z ≥ 3.1.

Next, we investigated the relationship between resulting clusters of task-based functional connectivity during the illusion condition and the susceptibility ratings. To do so, we extracted the connectivity values using a 5-mm radius sphere around the peak [right EBA connectivity clusters: right PPC (60,−20,44), right vlPFC (44,48,4) and SMA (4,28,50); left EBA connectivity: right PPC (60,−22,42), right vlPFC (40,48,12) and SMA (4,26,50)]. These connectivity values were correlated with susceptibility ratings, and significance was set at *p*<0.0083 (Bonferroni-corrected for six comparisons). The critical R^2^-value for a significance level set at *p*<0.0083 and n=18 was calculated to be r^2^=0.361.

## Results

### Behavioural Study

First, we sought to determine whether exposure to a finger-stretch illusion (see Fig 1a) in an MRI scanner with a different egocentric perspective to the usual direct and proximal perspective would affect the illusory experience. Thus, twelve healthy participants underwent the original finger-stretch illusion (with a proximal setup) and a modified finger-stretch illusion with distal setup akin to that used in the MRI environment (see Fig 1b). There were no significant differences in the experience of the two illusion setups (main effect of illusion: *F*_1,11_= 1.43, *p*= 0.2564, all post-hoc t-tests *p*> 0.05; Fig 1c and S1 Table). Therefore, the MRI setup was not significantly different than the proximal setup.

### Imaging Study

Eighteen participants experienced a finger-stretch illusion in a 3T MRI scanner. During the illusion, the participant’s left index finger was visually elongated with congruent tactile input. A control condition included all the same procedures without visual elongation. Participants provided trial-by-trial ratings of the extent to which they felt that their finger was actually being stretched in both conditions. Ratings were significantly greater during the illusion compared to the control condition (*p*< .0001), indicating that participants were susceptible to the illusion (Fig 2a). We created a susceptibility score based on the average difference scores between the illusion and control trials for each participant. Individual differences in behavioural measures have been shown to be reflected in brain structure^15^. To determine the structural gray matter underpinnings of individual differences in susceptibility, we performed a whole brain voxel-based morphometry (VBM) analysis. We found that bilateral extrastriate body area (EBA; part of the occipitotemporal cortex—OTC^16^) volumes were positively correlated with susceptibility (*r^2^*= 0.74; *p*_*FWE*_< 0.05, with a cluster-forming height threshold *p*< .001; Fig 2b-c, and S2 Table). In other words, the greater EBA volume, the more susceptible the participant was to the finger-stretch illusion.

**Fig 2.**
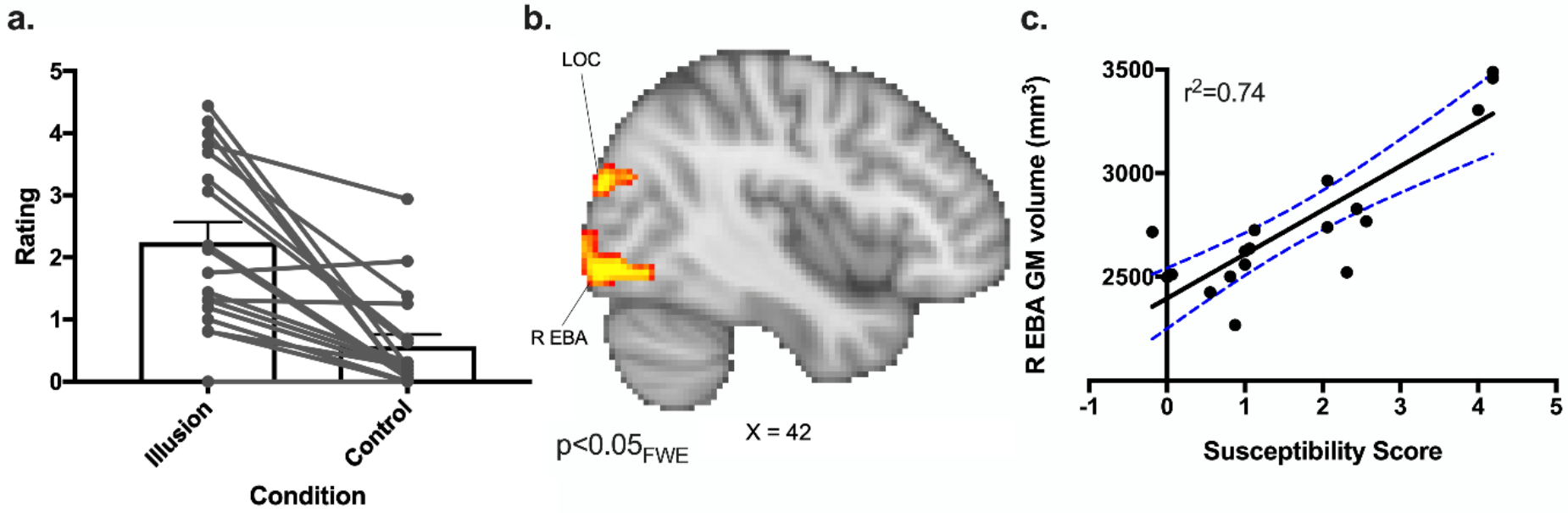
Susceptibility to illusion and neural correlates. (a) Individual participant ratings of the finger stretch illusion and control condition in the MRI scanner. These ratings (between 0 and 5) represent the susceptibility of the participants to the illusions. Mean (±SE) of ratings to the *statement “The extent to which you feel that your finger is actually being stretched’*” (n=18). Participant ratings were based on 16 trials of each condition. Ratings for the illusion were significantly higher than the control condition (p=0.00009; Cohen’s d= 1.19671). (b) Bilateral temporo-occipital gray matter volumes correlate with finger-stretch illusion susceptibility (the difference between the illusion and control conditions). Statistical images are cluster-corrected *p*<0.05 family-wise error, with a cluster-forming height threshold of p<.001. (c) Significant correlation of the right extrastriate body area gray matter volume with susceptibility ratings (r^2^=0.74). Note that both EBA were significantly correlated with susceptibility, and the right EBA is shown for simplicity. Blue lines represent 95% confidence intervals. Abbreviations: R EBA – right extrastriate body area; LOC – lateral occipital cortex.

### Whole brain activation during illusion

We determined whole brain activation in response to the finger-stretch illusion, compared to the control condition, and found that multisensory brain areas—the bilateral occipitotemporal junction, in the area of the EBA and fusiform body areas (FBA), the bilateral PPC, the bilateral lateral occipital cortex (LOC), and left PMv— showed greater activity during the illusion (Fig 3, S1 and S3 Figs, S3 and S4 Tables) [9, 17–19]. Notably, EBA did not show significant activation in the control condition (S1 and S3 Figs, and S5 Table). The activation of multisensory areas in response to the finger-stretch illusion is in line with other illusions [11,12].

**Fig 3.**
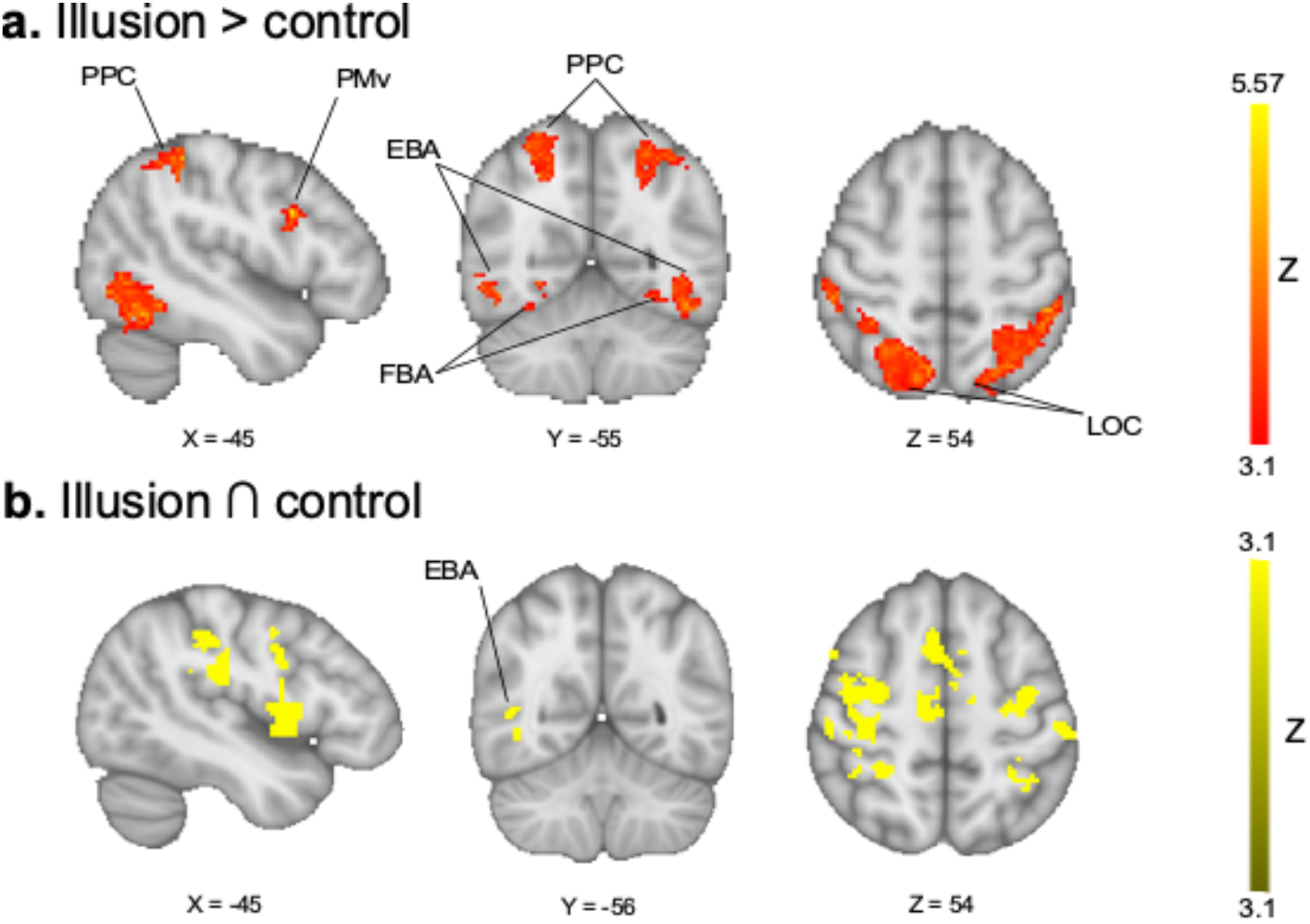
Contrast and conjunction analyses between finger-stretch and control illusion. (a) Brain activations in response to the finger-stretch illusion compared to a control illusion (n=18). (b) Conjunction map showing overlap of regions that show activation in both the finger-stretch illusion and the control illusion. Statistical images are cluster-corrected *p*_*FWE*_<0.05 (cluster-forming height threshold *Z*>3.1). Abbreviations: EBA – extrastriate body area; FBA – fusiform body area; LOC – lateral occipital cortex; PMv– ventral premotor cortex; PPC – posterior parietal cortex. Images are shown in radiological convention.

### Functional connectivity during illusion

Following our structural and functional brain imaging findings of a relationship between the EBA and bodily illusion (Figs 2 and 3), we performed a psychophysiological interaction (PPI) to determine which brain regions were functionally connected during the finger-stretch illusion. We found that the left and right EBA were functionally connected to the PPC, the supplementary motor area, and the lateral frontopolar cortex (*p*_*FWE*_<0.05, cluster-forming height threshold *Z*>3.1; Fig 4a, S4 Fig, and S6 and S7 Tables). Most notably, we report that the connectivity between the right EBA and the right PPC was significantly correlated to the strength of the illusion (r^2^=0.41, p=.0044; Fig 4b, S8 Table).

**Fig 4:**
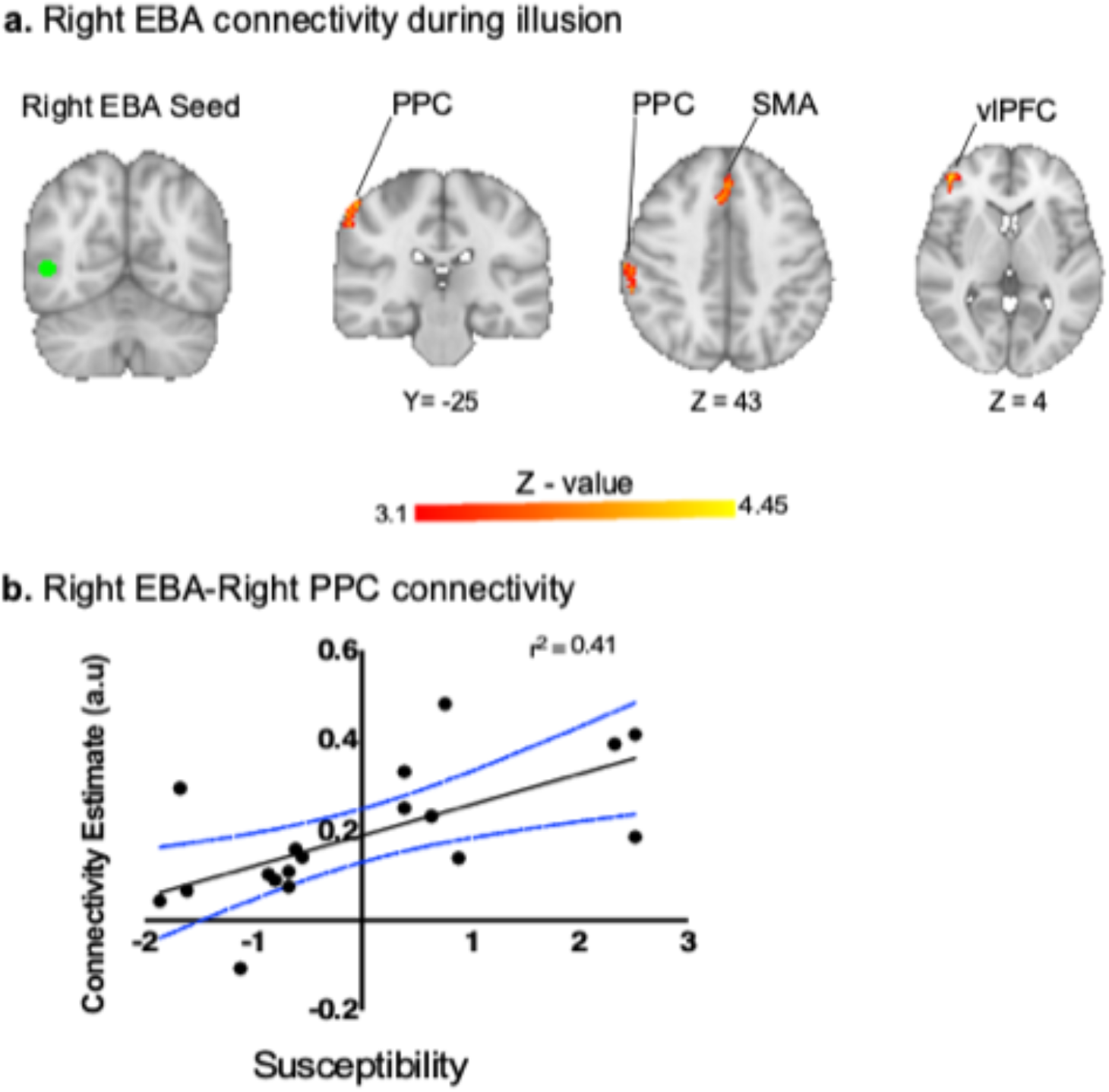
Functional connectivity during illusion with susceptibility scores. (a) Psychophysiological interaction of the right extrastriate body area (EBA; shown in green) during illusion condition. The EBA was significantly connected to the primary somatosensory cortex/posterior parietal cortex (S1/PPC), the supplementary motor area (SMA) and the ventrolateral prefrontal cortex (vlPFC). The statistical threshold includes cluster corrected threshold of *p*<.05 (*p*<.007, Bonferroni-corrected for PPIs performed) using a cluster forming threshold *Z*>3.1. Note that bilateral EBA showed functional connectivity to these regions, but the right is shown for simplicity, and because it is contralateral to the illusion. (b) Positive correlation between right EBA-PPC task-based functional connectivity during the illusion condition and susceptibility scores (r^2^=0.41, p<0.008; Bonferroni-corrected p<0.05 divided by six tests). Blue lines represent 95% confidence intervals. a.u.—arbitrary units. Images are shown in radiological convention.

## Discussion

We utilized a multimodal finger-stretch illusion to examine the behavioural and neural correlates of body image. We showed that the structure of the lateral occipitotemporal cortex, in the region of the EBA, was significantly correlated to how susceptible participants were to the illusion (Fig 2), and that multisensory regions were activated by the finger-stretch illusion (Fig 3), including bilateral lateral occipitotemporal cortex, in the region of the EBA (see S3 Fig 3), PPC and PMv. Finally, we show that the task-based functional connectivity of the occipitotemporal cortex—the putative EBA— with the PPC is significantly correlated with susceptibility scores (Fig 4). Together, these multimodal data indicate that the lateral occipitotemporal cortex, in the region of the putative EBA, encodes body image perception.

The EBA is an occipitotemporal region activated by congruent spatial and synchronous tactile stimuli [12,20]. The EBA has been shown to respond to various stimuli, including human touch, action, and motion [11, 21–23]. Additionally, the EBA is involved in visual perception and the processing of human body parts, and the body as a whole [16,24]. Previous studies have identified selective disruption of body perception upon interference by transcranial magnetic stimulation (TMS) to the EBA [25,26]. It is therefore expected that bilateral activation of the EBA would occur during the congruent visuo-tactile finger-stretch illusion and the control condition performed in the present study. The enhanced EBA activation in the illusion condition suggests that the EBA response is not only due to processing congruent visuo-tactile stimuli, but plays a direct role in body perception, more specifically in upper limb perception.

Enhanced PMv activity was observed during the illusion condition compared to the control condition. The PMv houses neurons with both visual and tactile receptive fields [27] and is important for the detection of visuo-proprioceptive congruence [12,28]. Our finding is in line with previous studies examining neural correlates of visual illusions, such as the rubber hand illusion (RHI) [29,30]. In addition, the PMv is thought to relate to higher-level processing of upper limb representation and the surrounding (peri-personal) space [10]. Studies have sought to elucidate the role of the PMv in the illusory experience and proprioception by utilizing a variety of tasks, such as arm positioning [12] and RHI manipulations [11,31]. The arm positioning task identifies brain regions active during congruent positioning of the unseen subject’s own arm and a virtual realistic arm^12^. The RHI paradigms manipulate either visual or tactile inputs, which control for the visualization of the rubber hand (somatic RHI) [31] and the interference of the experimenter (automated RHI) [11], respectively. The PMv responds to the somatic RHI [31] and arm positioning task [12], but not to the automated RHI [11], suggesting that this region is involved in processing tactile and proprioceptive inputs independent of human touch and action, as well as integrating congruent visual and proprioceptive information about arm position. The enhanced PMv activation observed in the present study therefore supports the role of the PMv in the integration of multisensory, visuotactile information in response to the finger-stretch illusion. This activation could support the role of the PMv in upper limb proprioception that is involved in body schema, by updating proprioceptive information upon receiving new visuotactile information.

Our second major finding shows significantly enhanced functional connectivity of the bilateral EBA to the PPC, supplementary motor area, and ventrolateral prefrontal cortex (vlPFC) during the finger-stretch illusion. Most notably, we report that the connectivity between the right EBA and the right PPC was significantly correlated to the strength of the illusion (r^2^=0.41, p=0.0044; Fig 3b). This finding is consistent with Limanowski & Blankenburg [20], who reported increased functional connectivity between the EBA and PPC during a visuo-proprioceptive RHI. However, the possibility of visual and proprioceptive integration occurring in the PPC rather than the EBA was not ruled out in their study. The PPC is thought to maintain a dynamic estimate of the perceptual representation of the body, in particular the hand region [32,33], that can be updated through multisensory integration [9,34]. Though other RHI studies have demonstrated increased EBA-PPC functional connectivity [29,35], they were unable to attribute it to the visuo-proprioceptive illusion. Rather than EBA-PPC connectivity, one RHI study demonstrated increased functional connectivity between the bilateral EBA and the primary somatosensory cortex (S1) [11]. Inhibitory stimulation of the S1 hand region by repetitive TMS in healthy participants resulted in an overestimation of the perceived size of their hand [36], suggesting that the S1 plays a role not only in somatosensation, but in the perception of body size—which is in line with the fine representation of the hands in S1, compared to coarser representations in higher order brain regions, such as the PPC [37]. A previous study proposed that the intraparietal sulcus (IPS), which separates the superior and inferior parietal lobules, minimizes mismatch between the incoming sensory information by integrating this visual information with tactile information. Specifically, the IPS integrates the somatosensory reference frame with the visual reference frame to minimize mismatch, and consequently increases its connectivity to the EBA [29]. Taken together, the connectivity of the EBA to other parietal multisensory regions underlies body shape and size during multisensory illusions. In line with these previous findings, we demonstrate that EBA-PPC functional connectivity is related to a person’s susceptibility to the finger-stretch illusion.

Information from multiple sensory channels are integrated in higher cognitive areas to construct a body representation [38]. Specifically, the convergent somatosensory, proprioceptive and visual inputs to the EBA are integrated and underlie human body shape perception [16,39]. Our data suggest that the structure and functional connectivity of the EBA not only encodes the shape of body parts, but also how susceptible a participant is to disturbances of shape. Our structural and functional connectivity findings are the first evidence to demonstrate the role of EBA in perceiving changes to body shape. More importantly, the functional connectivity between EBA-PPC is predictive of how susceptible the participant is to a change in this body shape. Disturbances of body image—i.e., body perception disturbances—have been reported in multiple disorders [1], including anorexia nervosa, bulimia nervosa, chronic pain [40–42] and somatoparaphrenia [43]. Despite the prevalence of these disturbances across various disorders, the neural underpinnings remain unknown. Identifying these mechanisms can provide novel therapeutic targets for these disorders.

## Acknowledgements

This study was entirely funded by a University of the West of England QR grant number 201415/31.

## Supporting information

### Supporting Methods

#### Functional localizer of EBA

The EBA is a functionally distinct region. To ensure brain activations in the lateral occipital cortex were consistent with the EBA, we used a functional localizer derived from meta-analyses that used the term “body” in Neurosynth (www.neurosynth.org). We found 552 studies with 19,337 activations that produced an association test map displaying the bilateral EBA. The association test map demonstrates that there is an association between voxels in the map and studies that use the term “body” in their abstracts^48^. We compared the association test map with our illusion > control contrast to identify overlapping voxels. Images were cluster-corrected *pFDR*<0.01 for the association map, and *p*_*FWE*_<0.05 for the contrast, thresholded at *Z* >3.1 and binarized (S2 Fig).

### Supporting Results

**Supporting Fig 1.**
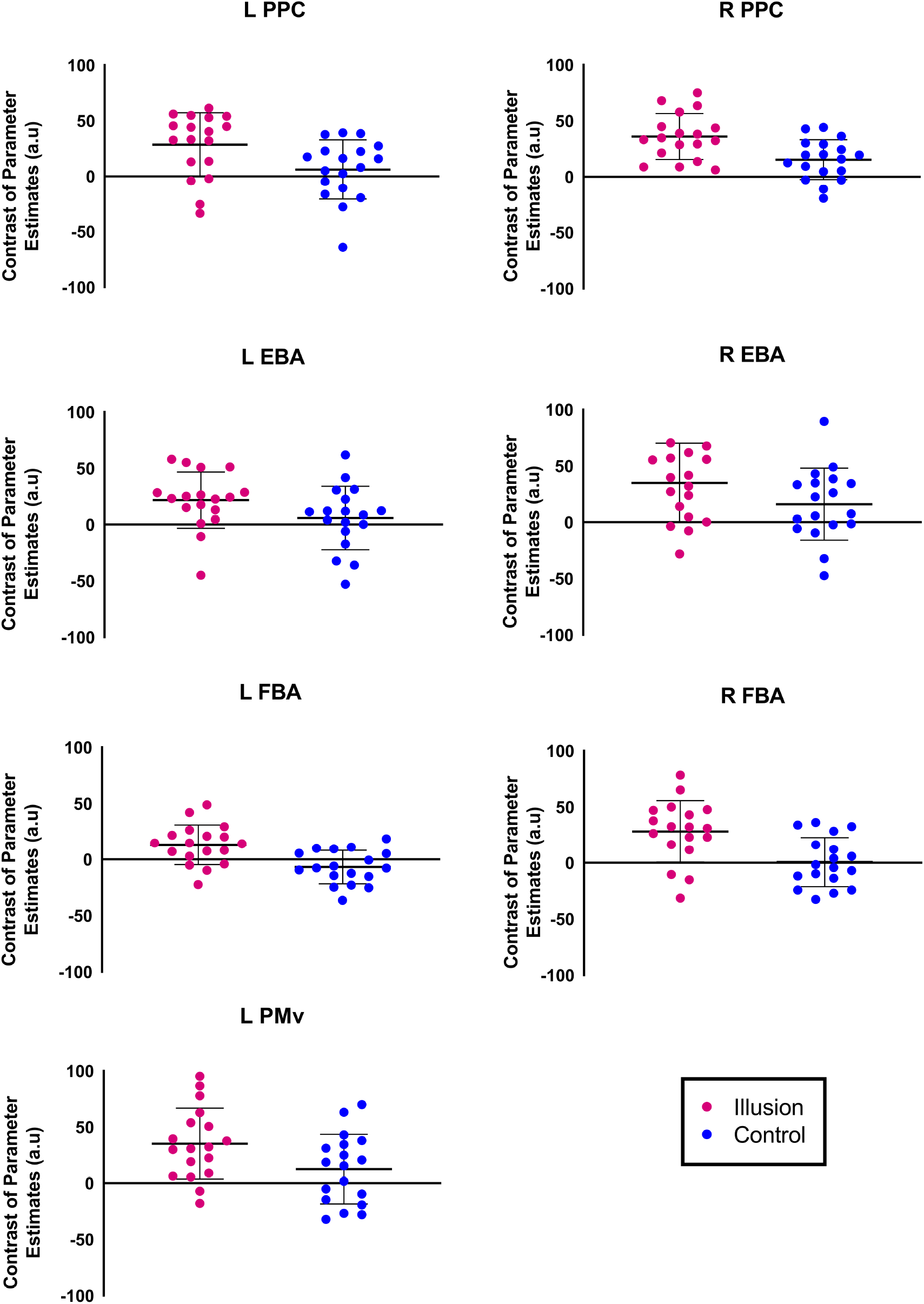
Activation differences between the illusion and control conditions. Mean and standard error of BOLD activity for regions with significantly greater activation in the illusion condition (red) compared to the control condition (blue; cluster corrected p<0.05, with a cluster-forming height threshold of Z>3.1). Abbreviations: a.u – arbitrary unit; L – left; R – right; EBA – extrastriate body area; FBA – fusiform body area; PMv – ventral premotor cortex; PPC – posterior parietal cortex.

**Supporting Fig 2.**
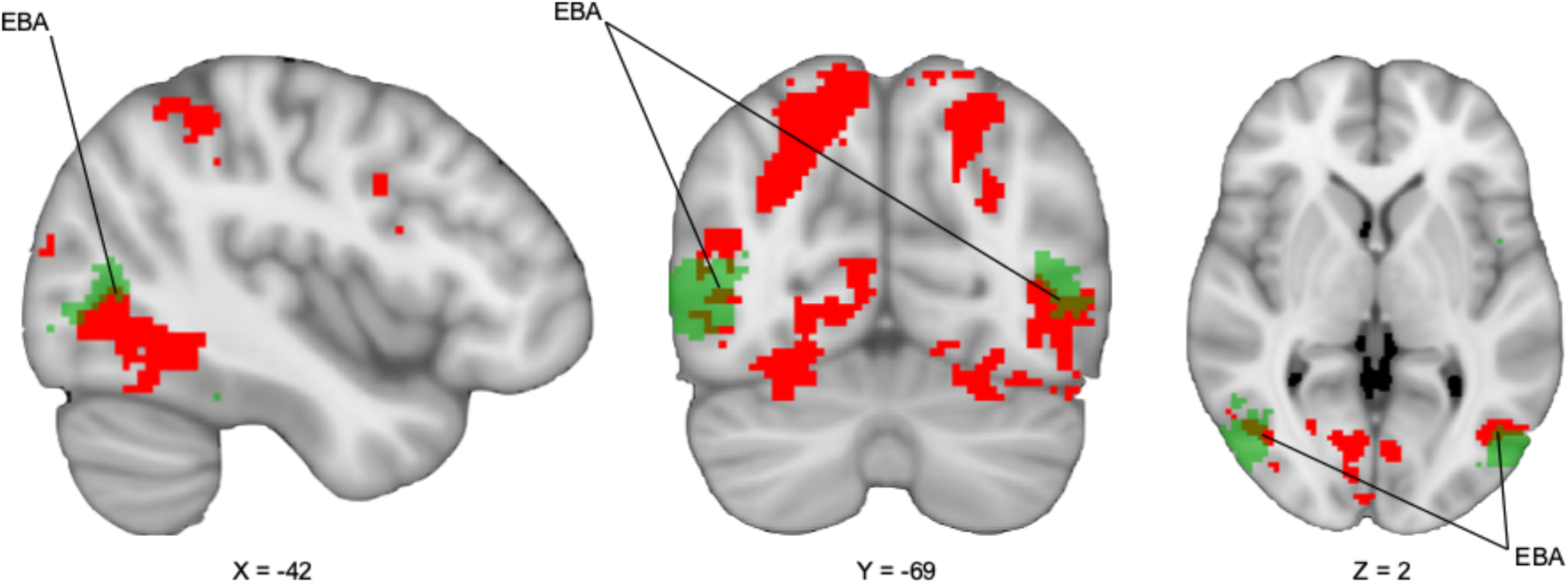
Extrastriate body area functional localizer. A meta-analysis was performed on neuroimaging studies using the Neurosynth database. We identified voxels associated with the keyword “body” across the database. The resulting map shows the EBA (green; cluster-corrected *p*_*FDR*_<0.01) overlaid on our contrast map of the illusion condition versus the control condition (red; cluster-corrected *p*_*FWE*_<0.05). Thresholded images were binarized for visualization purposes. Abbreviations: EBA – extrastriate body area. Images are shown in radiological convention.

**Supporting Fig 3.**
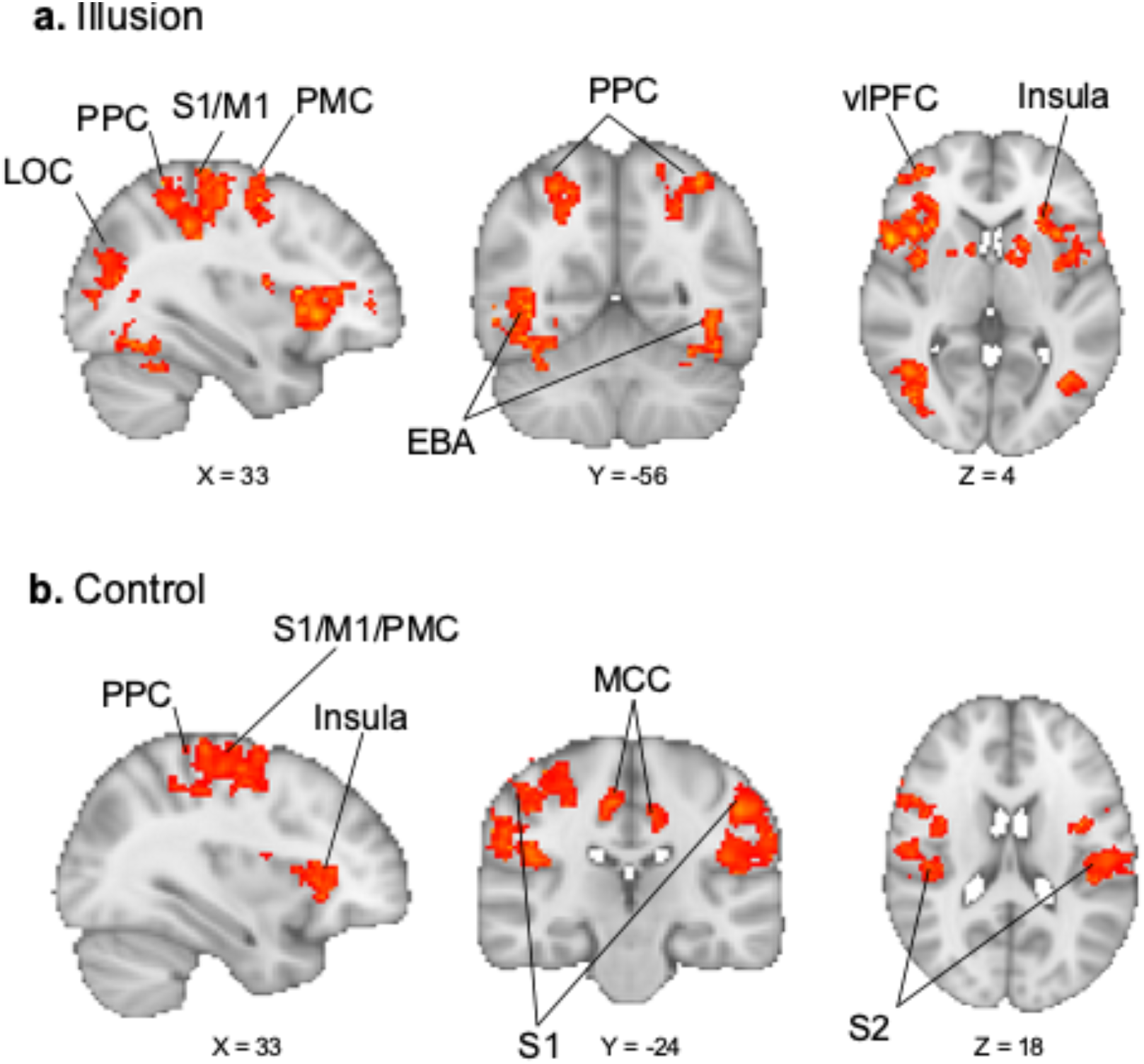
Brain activation associated with illusion and control. (a) mean activation for finger-stretch illusion task (n=18). (b) mean activation for control illusion (n=18). Statistical images are cluster-corrected *p*_*FWE*_<0.05 (cluster-forming height threshold *Z*>3.1). All the illustrated activation clusters are shown on an MNI 152 2 mm brain mask in standard space. Abbreviations: LOC – lateral occipital cortex; PPC – posterior parietal cortex; S1-the primary somatosensory cortex; S2 – secondary somatosensory cortex; M1 – primary motor cortex; PMC – premotor cortex; EBA – extrastriate body area; vlPFC – ventrolateral prefrontal cortex; MCC – middle cingulate cortex; OTC – occipitotemporal cortex. Images are shown in radiological convention.

**Supporting Fig 4.**
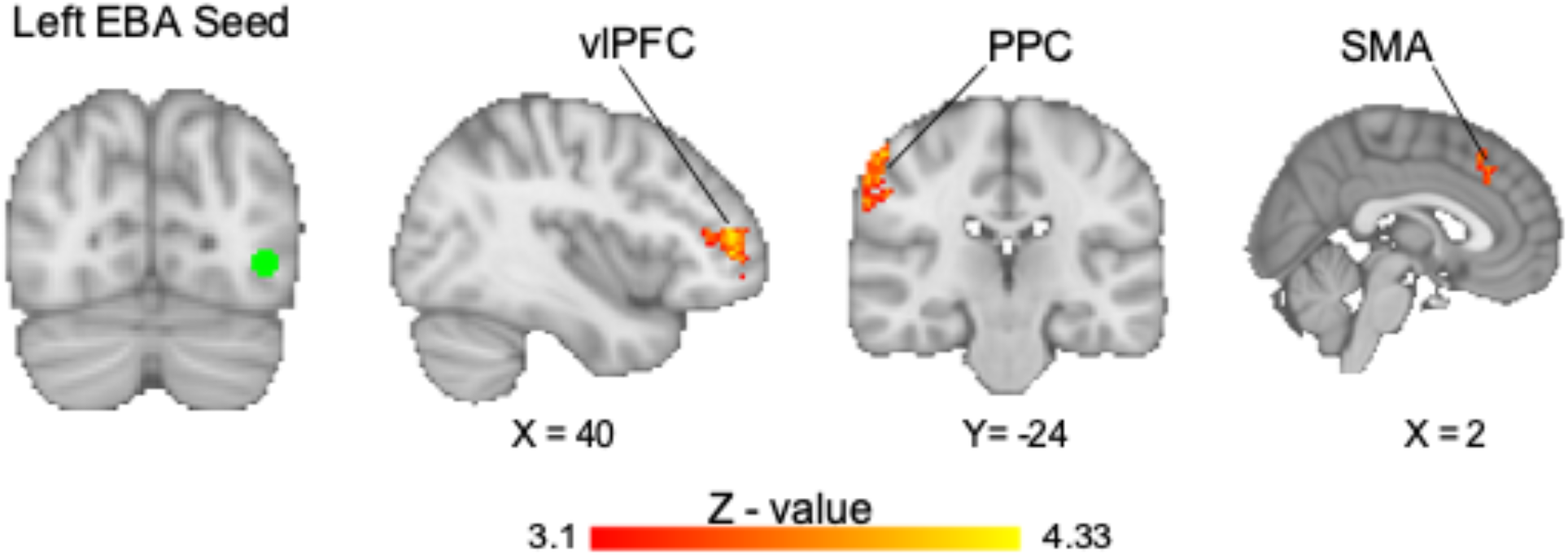
Task-based functional connectivity of the left extrastriate body area (EBA). A psychophysiological interaction of the left extrastriate body area (EBA; shown in green) was performed during illusion condition. The EBA was significantly connected to the primary somatosensory cortex/posterior parietal cortex (S1/PPC), the supplementary motor area (SMA) and the ventrolateral prefrontal cortex (vlPFC). The statistical threshold includes cluster corrected threshold of *p*<.05 (*p*<.007, Bonferroni-corrected for PPIs performed) using a cluster forming threshold *Z*>3.1. Images are shown in radiological convention.

**Supporting Table 1.** Behavioural Study post-hoc paired t-tests. The t-tests compare behavioural study statement ratings about the finger stretch illusion in the proximal and distal setups.

| Statement | Mean Difference | Standard Error of the Mean | T-value | Degrees of Freedom | p-value |
| --- | --- | --- | --- | --- | --- |
| It felt like my finger was really being stretched | -0.38 | 0.255 | -1.47 | 11 | 0.169 |
| I feel like my finger is longer than normal | 0.46 | 0.428 | 1.07 | 11 | 0.308 |
| I feel like I am watching myself | 0.25 | 0.392 | 0.64 | 11 | 0.536 |
| I feel like the finger I'm seeing belongs to me | 0.29 | 0.406 | 0.72 | 11 | 0.487 |
| My finger was getting hot | 0.71 | 0.377 | 1.88 | 11 | 0.087 |
| It felt like I had 2 left index fingers | 0.25 | 0.334 | 0.75 | 11 | 0.47 |

**Supporting Table 2.** Regions with gray matter volume correlated to finger-stretch illusion susceptibility scores

| Regions | Cluster extent (mm<sup>3</sup>) | Peak MNI |  |  | T-value |
| --- | --- | --- | --- | --- | --- |
|  |  | X | Y | Z |  |
| Right Occipitotemporal cortex (EBA) |  | 45 | -81 | -10 | 6.47 |
| Right Occipitotemporal cortex | 2557 | 48 | -87 | 3 | 5.50 |
| Right Superior Lateral Occipital Cortex |  | 44 | -84 | 21 | 5.23 |
| Left Occipitotemporal cortex (EBA) |  | -36 | -92 | 27 | 6.20 |
| Left Occipitotemporal cortex | 1645 | -46 | -88 | 8 | 4.74 |
| Left Superior Lateral Occipital Cortex |  | -39 | -94 | 8 | 4.39 |

Abbreviations: EBA – Extrastriate Body Area; FWE – Family-wise error; MNI – Montreal Neurological Institute.

**Supporting Table 3.** Brain regions with increased BOLD activity during illusion compared to control tasks.

| Regions | Cluster extent<br>(number of voxels) | Peak MNI |  |  | Z-score |
| --- | --- | --- | --- | --- | --- |
|  |  | X | Y | Z |  |
| Right Lateral Occipital Cortex (superior division) | 5285 | 30 | -84 | 34 | 5.07 |
| Right Lateral Occipital Cortex (superior division) |  | 34 | -78 | 24 | 4.99 |
| Right Superior Parietal Lobule |  | 14 | -72 | 62 | 4.99 |
| Right Superior Parietal Lobule |  | 14 | -78 | 50 | 4.99 |
| Right Extrastriate Body Area |  | 12 | -74 | -10 | 4.94 |
| Right Inferior Parietal Lobule |  | 38 | -80 | 24 | 4.93 |
| Left Posterior Parietal cortex | 1092 | -46 | -40 | 54 | 4.77 |
| Left Posterior Parietal cortex |  | -26 | -66 | 50 | 4.68 |
| Left Posterior Parietal cortex |  | -32 | -62 | 58 | 4.61 |
| Left Posterior Parietal cortex |  | -30 | -62 | 50 | 4.56 |
| Left Posterior Parietal cortex |  | -18 | -70 | 52 | 4.47 |
| Left Extrastriate Body Area | 692 | -42 | -74 | -2 | 4.79 |
| Left Lateral Occipitotemporal Sulcus |  | -46 | -58 | -16 | 4.74 |
| Left Extrastriate Body Area |  | -52 | -64 | 0 | 4.44 |
| Left Lateral Occipitotemporal Sulcus |  | -40 | -68 | -6 | 4.43 |
| Left Occipitotemporal Area |  | -52 | -62 | -14 | 4.42 |
| Left Occipitotemporal Area |  | -54 | -68 | 0 | 4.40 |
| Left Posterior Parietal cortex |  | -44 | -46 | 54 | 4.43 |
| Left Lateral Occipital Cortex (superior division) | 211 | -40 | -84 | 16 | 4.56 |
| Left Inferior Parietal Sulcus |  | -30 | -70 | 32 | 4.45 |
| Left Lateral Occipital Cortex (superior division) |  | -40 | -84 | 20 | 4.31 |
| Left Lateral Occipital Cortex (superior division) |  | -36 | -86 | 26 | 4.11 |
| Left Lateral Occipital Cortex (superior division) |  | -32 | -80 | 34 | 4.06 |
| Left Lateral Occipital Cortex (superior division) |  | -26 | -68 | 26 | 3.84 |
| Left Premotor cortex | 85 | -46 | 10 | 30 | 5.52 |
| Left Premotor cortex |  | -40 | 2 | 34 | 4.01 |
| Left Premotor cortex |  | -52 | 10 | 24 | 3.85 |

Abbreviations: MNI – Montreal Neurological Institute. The statistical threshold was set to a cluster-corrected $p < 0.05$ , with a cluster-forming height threshold of $Z > 3.1$ .

**Supporting Table 4.** Brain regions with increased BOLD activity during illusion task.

| Regions | Cluster Extent<br>(number of voxels) | Peak MNI |  |  | Z score |
| --- | --- | --- | --- | --- | --- |
|  |  | X | Y | Z |  |
| Right Lateral Occipitotemporal Sulcus | 12196 | 46 | -64 | 8 | 6.91 |
| Right Dorsolateral Prefrontal Cortex |  | 42 | 12 | 36 | 6.63 |
| Right Fusiform Gyrus |  | 30 | -64 | -16 | 6.59 |
| Right Inferior Frontal Gyrus |  | 56 | 8 | 26 | 6.55 |
| Right Dorsolateral Prefrontal Cortex |  | 58 | 12 | 36 | 6.35 |
| Right Inferior Parietal Lobule |  | 60 | -32 | 30 | 6.25 |
| Left Postcentral Gyrus | 2683 | -50 | -26 | 40 | 5.34 |
| Left Primary Somatosensory Cortex |  | -54 | -30 | 52 | 5.21 |
| Left Supramarginal Gyrus |  | -50 | -26 | 36 | 5.08 |
| Left Posterior Parietal Cortex |  | -36 | -46 | 52 | 5.08 |
| Left Posterior Parietal Cortex |  | -36 | -52 | 54 | 4.92 |
| Left Supramarginal Gyrus |  | -54 | -24 | 46 | 4.77 |
| Left Premotor Cortex | 2035 | -54 | 6 | 30 | 5.76 |
| Left Premotor Cortex |  | -40 | 0 | 36 | 5.20 |
| Left Dorsal Anterior Insula |  | -32 | 18 | 8 | 5.16 |
| Left Anterior Insula |  | -32 | 26 | -4 | 5.05 |
| Left Premotor Cortex |  | -46 | 2 | 34 | 4.77 |
| Left Premotor Cortex |  | -50 | 4 | 34 | 4.75 |
| Left Occipitotemporal Cortex (EBA) | 1022 | -42 | -74 | -8 | 4.83 |
| Left Occipitotemporal Cortex |  | -42 | -62 | -6 | 4.73 |
| Left Occipitotemporal Cortex |  | -40 | -62 | -2 | 4.65 |
| Left Occipitotemporal Cortex |  | -42 | -64 | -12 | 4.51 |
| Left Thalamus | 220 | -18 | 12 | 2 | 4.38 |
| Left Putamen |  | -18 | 8 | 2 | 4.30 |
| Left Caudate |  | -14 | 10 | 0 | 4.25 |
| Left Thalamus |  | -10 | 0 | 0 | 4.12 |
| Left Caudate |  | -12 | 4 | 4 | 4.01 |
| Left Caudate |  | -14 | 14 | -2 | 3.94 |
| Left Prefrontal Cortex – Anterior Middle Frontal Gyrus | 154 | -48 | 40 | 12 | 4.70 |
| Left Ventrolateral Prefrontal Cortex |  | -40 | 40 | 14 | 4.56 |
| Left Ventrolateral Prefrontal Cortex |  | -44 | 40 | 20 | 4.15 |
| Left Ventrolateral Prefrontal Cortex |  | -36 | 46 | 12 | 3.97 |
| Left Ventrolateral Prefrontal Cortex |  | -40 | 48 | 14 | 3.77 |
| Left Ventrolateral Prefrontal Cortex |  | -44 | 46 | 14 | 3.69 |
| Right Caudate | 123 | 18 | 4 | 12 | 4.09 |
| Right Putamen |  | 20 | 8 | 10 | 4.03 |
| Right Caudate |  | 12 | 8 | 8 | 4.00 |
| Right Putamen |  | 22 | 6 | 6 | 3.99 |
| Right Caudate |  | 12 | 6 | 2 | 3.90 |
| Right Caudate |  | 16 | 4 | 8 | 3.72 |
| Right Lateral Orbitofrontal Cortex | 81 | 26 | 44 | -18 | 4.58 |
| Right Lateral Orbitofrontal Cortex |  | 30 | 44 | -14 | 4.49 |
| Right Lateral Orbitofrontal Cortex |  | 26 | 42 | -12 | 4.27 |
| Right Lateral Orbitofrontal Cortex |  | 22 | 40 | -14 | 4.19 |
| Right Lateral Orbitofrontal Cortex |  | 16 | 50 | -20 | 3.27 |

Abbreviations: MNI – Montreal Neurological Institute. The statistical threshold was set to a cluster-corrected $p < 0.05$ , with a cluster-forming height threshold of $Z > 3.1$ .

**Supporting Table 5.**
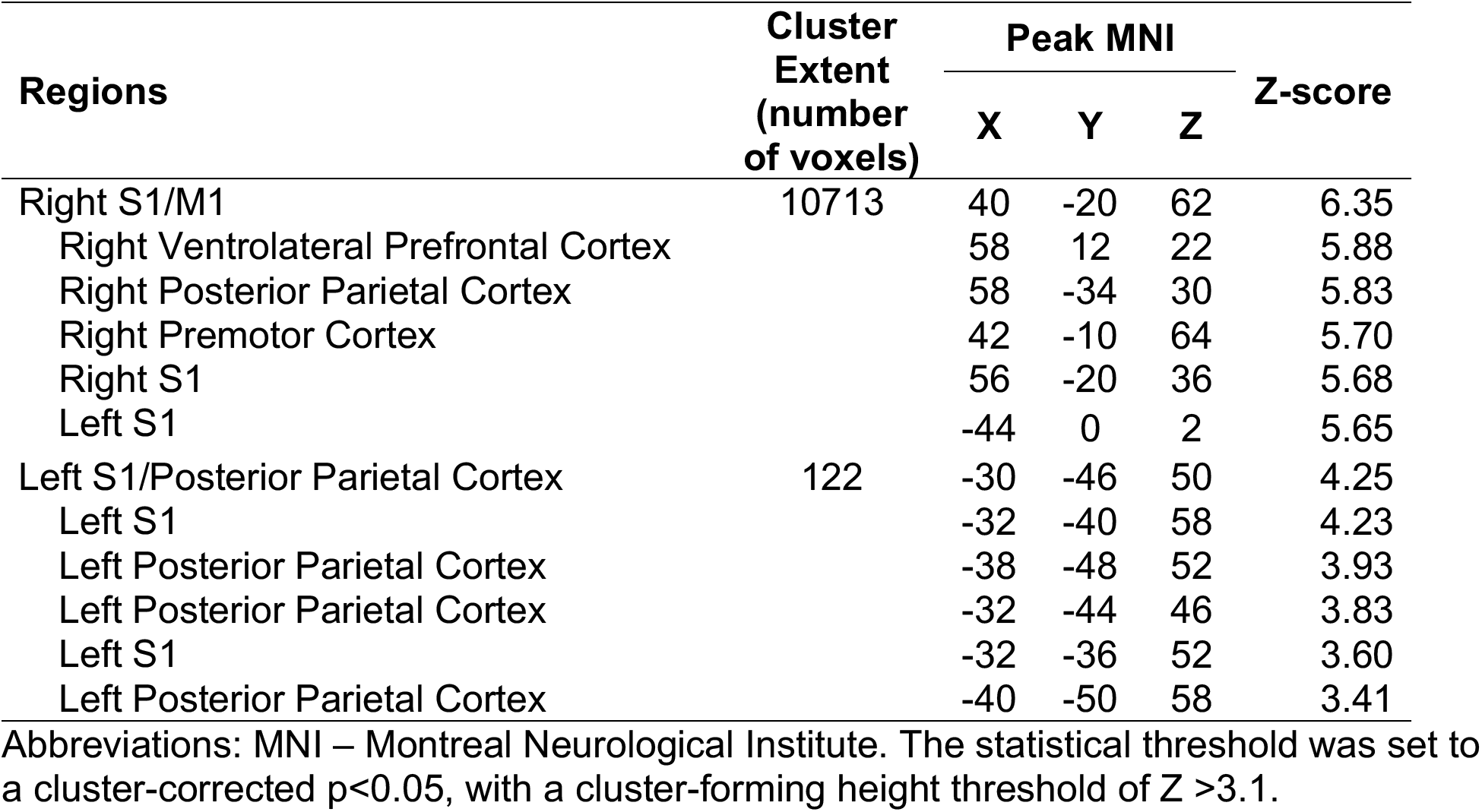
Brain regions with increased BOLD activity during control task.

**Supporting Table 6.** Brain regions with stronger task-based functional connectivity to the right extrastriate body area during the finger-stretch illusion.

| Regions | Cluster extent<br>(number of voxels) | Peak MNI |  |  | Z-score |
| --- | --- | --- | --- | --- | --- |
|  |  | X | Y | Z |  |
| Right Ventrolateral Prefrontal cortex | 306 | 44 | 48 | 4 | 4.23 |
| Right Ventrolateral Prefrontal cortex |  | 34 | 46 | 10 | 4.02 |
| Right Ventrolateral Prefrontal cortex |  | 46 | 28 | 12 | 3.99 |
| Right Ventrolateral Prefrontal cortex |  | 40 | 48 | 10 | 3.99 |
| Right Ventrolateral Prefrontal cortex |  | 42 | 38 | 10 | 3.96 |
| Right Ventrolateral Prefrontal cortex |  | 46 | 24 | 12 | 3.76 |
| Right Posterior Parietal Cortex/S1 | 213 | 60 | -20 | 48 | 4.27 |
| Right Posterior Parietal Cortex/S1 |  | 64 | -20 | 44 | 4.16 |
| Right Posterior Parietal Cortex |  | 56 | -26 | 56 | 4.16 |
| Right Inferior Parietal Lobule |  | 60 | -24 | 44 | 4.15 |
| Right Inferior Parietal Lobule |  | 58 | -30 | 46 | 4.01 |
| Right Inferior Parietal Lobule |  | 62 | -30 | 46 | 3.98 |
| Right Supplementary Motor Area | 152 | 4 | 28 | 50 | 4.41 |
| Supplementary Motor Area |  | 0 | 30 | 40 | 4.33 |
| Supplementary Motor Area |  | 0 | 30 | 34 | 4.24 |
| Right Supplementary Motor Area/MCC |  | 6 | 22 | 42 | 3.91 |
| Supplementary Motor Area/MCC |  | 0 | 24 | 36 | 3.88 |

Abbreviations: aMCC – anterior midcingulate cortex; MNI – Montreal Neurological Institute; S1 – Primary somatosensory cortex. The statistical threshold for was set to $p < 0.05$ ( $p \leq 0.007$ , corrected for multiple comparisons), with a cluster-forming height threshold $Z \geq 3.1$ .

**Supplementary Table 7.** Brain regions with stronger task-based functional connectivity to the left extrastriate body area during the finger-stretch illusion.

| Regions | Cluster Extent<br>(number of voxels) | Peak MNI |  |  | Z-score |
| --- | --- | --- | --- | --- | --- |
|  |  | X | Y | Z |  |
| Right Inferior Parietal Lobule | 285 | 60 | -22 | 42 | 4.21 |
| Right Inferior Parietal Lobule |  | 58 | -30 | 50 | 4.15 |
| Right S1 |  | 56 | -24 | 54 | 4.13 |
| Right Inferior Parietal Lobule |  | 56 | -34 | 52 | 4.04 |
| Right S1/ Inferior Parietal Lobule |  | 62 | -22 | 46 | 4.00 |
| Right Supramarginal Gyrus |  | 64 | -24 | 34 | 3.91 |
| Right Ventrolateral Prefrontal cortex | 243 | 40 | 48 | 12 | 4.29 |
| Right Ventrolateral Prefrontal cortex |  | 40 | 48 | 4 | 4.25 |
| Right Ventrolateral Prefrontal cortex |  | 46 | 50 | -4 | 3.91 |
| Right Ventrolateral Prefrontal cortex |  | 46 | 38 | 18 | 3.84 |
| Right Ventrolateral Prefrontal cortex |  | 46 | 46 | 10 | 3.74 |
| Right Supplementary Motor Area | 102 | 4 | 26 | 50 | 4.14 |
| Supplementary Motor Area |  | 0 | 34 | 42 | 3.82 |
| Right aMCC |  | 6 | 26 | 42 | 3.78 |
| Right Supplementary Motor Area |  | 0 | 28 | 34 | 3.75 |

Abbreviations: aMCC – anterior midcingulate cortex; MNI – Montreal Neurological Institute; S1 – Primary somatosensory cortex. The statistical threshold for was set to $p < 0.05$ ( $p \leq 0.007$ , corrected for multiple comparisons), with a cluster-forming height threshold $Z \geq 3.1$ .

**Supporting Table 8.** R-squared (R^2^) values of the relationship between task-based functional connectivity of the EBA during the illusion task and susceptibility scores.

|  | SMA | Right vIPFC | Right PPC |
| --- | --- | --- | --- |
| Left EBA | 0.13 | 0.08 | 0.23 |
| Right EBA | 0.18 | 0.09 | **0.41** |

The critical $R^2$ value for a significance value of $p < 0.0083$ (Bonferroni-corrected for multiple comparisons) and $n=18$ is $R^2=0.36$ . **Bold** font indicates a significant correlation. Abbreviations: EBA – extrastriate body area; SMA – supplementary motor area; vIPFC – ventrolateral prefrontal cortex; PPC – posterior parietal cortex.

